# Building a sequence map of the pig pan-genome from multiple *de novo* assemblies and Hi-C data

**DOI:** 10.1101/459453

**Authors:** Xiaomeng Tian, Ran Li, Weiwei Fu, Yan Li, Xihong Wang, Ming Li, Duo Du, Qianzi Tang, Yudong Cai, Yiming Long, Yue Zhao, Mingzhou Li, Yu Jiang

**Affiliations:** Key Laboratory of Animal Genetics, Breeding and Reproduction of Shaanxi Province, College of Animal Science and Technology, Northwest A&F University, Yangling 712100, China.; College of Animal Science and Technology, Sichuan Agricultural University, Chengdu 611130, China.

## Abstract

Pigs (*Sus scrofa*) exhibit diverse phenotypes in different breeds shaped by the combined effects of various local adaptation and artificial selection. To comprehensively characterize the genetic diversity of pigs, we construct a pig pan-genome by comparing genome assemblies of 11 representative pig breeds with the reference genome (Sscrofa11.1). Approximately 72.5 Mb non-redundant sequences were identified as pan-sequences which were absent from the Sscrofa11.1. On average, 41.7 kb of spurious heterozygous SNPs per individual are removed and 12.9 kb novel SNPs per individual are recovered using pan-genome as the reference for SNP calling, thereby providing enhanced resolution for genetic diversity in pigs. Homolog annotation and analysis using RNA-seq and Hi-C data indicate that these pan-sequences contain protein-coding regions and regulatory elements. These pan-sequences can further improve the interpretation of local 3D structure. The pan-genome as well as the accompanied web-based database will serve as a primary resource for exploration of genetic diversity and promote pig breeding and biomedical research.

## Introduction

*Sus scrofa* (i.e., pig or swine) is of enormous agricultural significance and is an attractive biomedical model. Pigs were domesticated from wild boars independently in Anatolia and East Asia approximately 10,000 years ago following long-term gene flow from their local wild counterparts (Larson, et al. 2005; Groenen, et al. 2012; Frantz, et al. 2015). The combined effects of local adaptation and human-driven artificial selection have shaped the genomic diversity of pigs and form the present various phenotypes. However, to date, these variations have been largely interpreted in the context of the annotated representative reference genome by aligning short reads to it. Increasing evidence suggests that a single individual genome is insufficient to capture all genetic diversities within a species since reference genome may have gaps, mis-assigned regions, or unable to provide a repository for all sequences (Golicz, Batley, et al. 2016). Alternatively, comparisons of independently *de novo*-assembled genomes and a reference genome sequence promise a more accurate and comprehensive understanding of genetic variations within a species (Li, et al. 2014; Schatz, et al. 2014).

Most recently, the pan-genome, the non-redundant collection of all genomic sequences of a species, has emerged as a fundamental resource for unlocking natural diversity in eukaryotes. Intraspecific comparisons in plants (e.g., soybean (Li, et al. 2014), Brassica oleracea (Golicz, Bayer, et al. 2016), Brachypodium distachyon (Gordon, et al. 2017) and rice (Zhao, et al. 2018)) and in animals (e.g., mosquitoes (Neafsey, et al. 2015), macaques (Yan, et al. 2011) and humans (Li, et al. 2010; Maretty, et al. 2017)) have revealed surprisingly large amounts of variation within a species. To build a high-quality pan-genome, a number of individual genomes are required (Li, et al. 2014; Monat, et al. 2017; Wong, et al. 2018), which remains an obstacle for most mammalian species. The current pig genome (Sscrofa11.1) represents one of the most continuous assemblies in mammalian species (**Supplementary Fig. 1**) and is from one individual (Duroc breed). In addition, our previous studies generated *de novo* assemblies of ten geographically and phenotypically representative pig genomes from Eurasia (Li, et al. 2013; Li, et al. 2017). Together with the assembly of Chinese Wuzhishan boar (Fang, et al. 2012), the availability of 12 pig genomes has provided an unprecedented opportunity to investigate their genetic differences at the individual, ethnic/breed or continental level.

Here we carried out an in-depth comparison between 11 *de novo* assemblies and the reference genome by analysis of the assembly-versus-assembly alignment.

The final pan-genome comprises 39,744 (total length: 72.5 Mb) newly added sequences and of which 607 demonstrate coding potential. Furthermore, the three-dimensional (3D) spatial structure of pan-genome was depicted by revealing the characteristics of pan-genome in A/B compartment (generally euchromatic and heterochromatic regions) and topologically associating domain (TAD). We also build a pig pan-genome database (PIGPAN, http://animal.nwsuaf.edu.cn/code/index.php/panPig) which can serve as a fundamental resource for unlocking variations within diverse pig breeds.

## Results

### Initial characterization of pan-sequences in the pig genome

To construct the pig pan-genome, we first aligned 11 assemblies from 11 genetically distinct breeds (five from Europe, and six from China) against Sscrofa11.1 using BLASTN to generate the unaligned sequences (**Fig. 1a and Supplementary Fig. 2**). The length of the unaligned sequences in the Chinese pigs was significantly longer than those in the European pigs (P <0.01) since the reference genome is from a European pig (**Fig. 1a**). As expected, the Wuzhishan assembly had the largest number of sequences because this sample is the only male individual among the 11 assemblies and can provide many male-specific sequences (**Fig. 1a and Supplementary Table 2**). After removing redundant sequences, we obtained 39,744 sequences with a total length of 72.5 Mb (**Fig. 1b**), which were absent from Sscrofa11.1 and thus were defined as pan-sequences. The content of the repetitive elements (45.91%) and GC (44.61%) in these sequences were slightly higher than those in Sscrofa11.1 (45.19% and 41.5%, respectively) (**Fig. 1a and Supplementary Table 3**). Notably, 44% (32 Mb) of the pan-sequences can be assigned to a unique assembly, highlighting the limitations of using one single assembly (**Fig. 1b**). All of the pan-sequences were longer than 300 bp. Among them, pan-sequences that are over 5 kb contributed 57% of the total length (**Fig. 1c**).

**Fig. 1.**
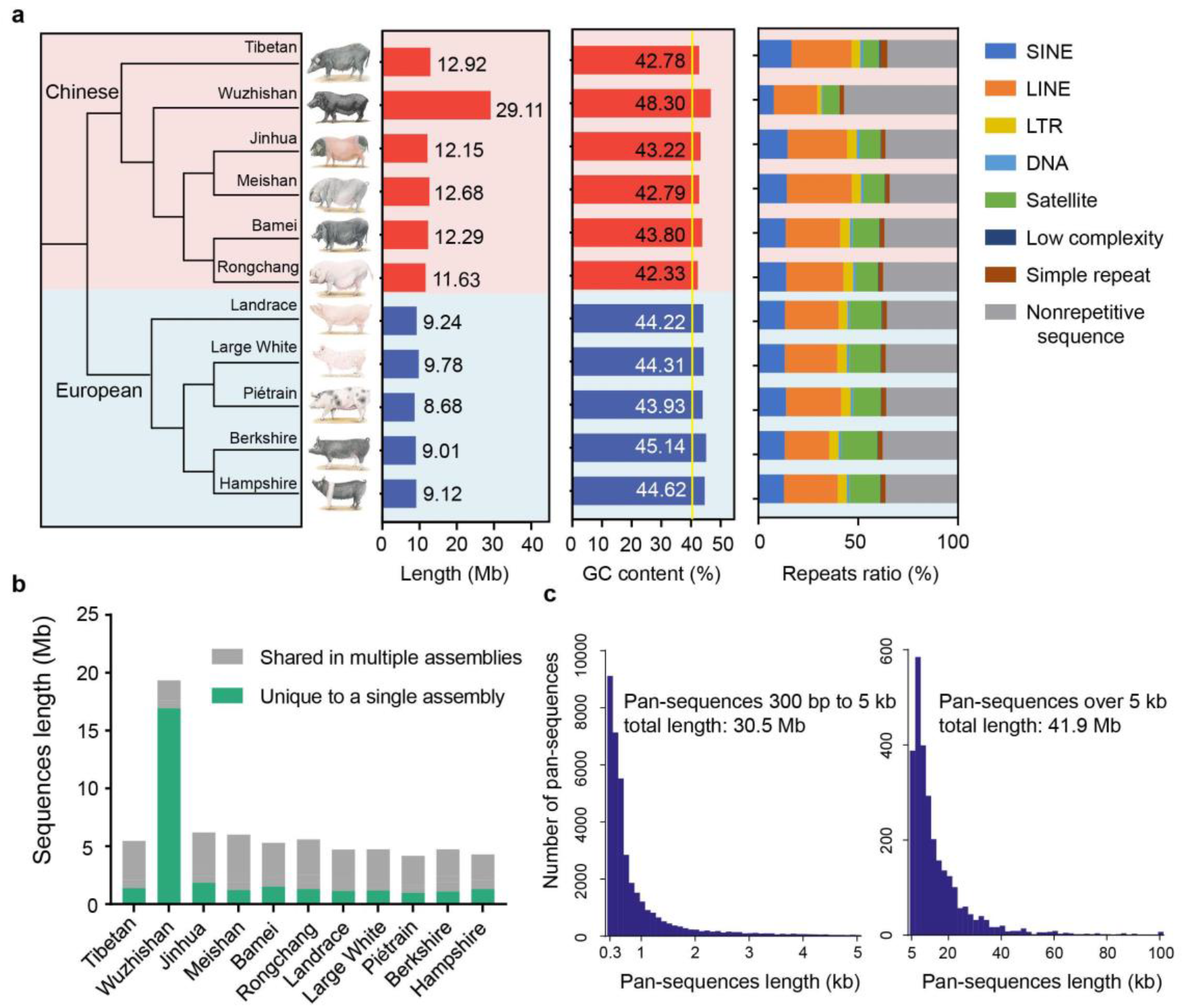
Construction of the pig pan-genome and the characterization of pan-sequences. **a** Maximum likelihood phylogenetic tree, sequence length, GC content and repeat composition of missing sequences identified in each individual assembly of eleven breeds (left to right). **b** The total sequence length and breed-specific sequence length of each breed for non-redundant pan-sequences. **c** Length distribution of all pan-sequences. (Wuzhishan pigs had the largest number of sequences because this individual is the only male among all the 11 assemblies and the sequencing platform of this individual differed from that used for other samples).

To validate the authenticity of the pan-sequences, we first compared these sequences to ten mammalian genomes and found that the majority (67.5%) of the pan-sequences had homologs in these genomes (*E*-value < 1e-5) (**Fig. 2a and Supplementary Table 4**). As expected, Cetacea has the greatest number of best hits in accordance with their close evolutionary relationship with pig (**Fig. 2a**). To explore the potential presence or absence of protein-coding genes within the pan-sequences, we aligned these pan-sequences to Refseq proteins from pig, cattle, goat, human, sheep and sperm whale using TBLASTN (*E*-value < 1e-5). The most significantly overrepresented hits were members of the olfactory receptor family (12.4%), which is the largest gene superfamily in vertebrates (Zhang and Firestein 2002) followed by other highly variable families (typically, O-methyltransferase domain-containing proteins), showing that these additional sequences are more variable than the reference set (**Fig. 2b**). Especially, 18 new and complete olfactory receptor genes were identified (**Supplementary Fig. 3**), implying that our pan-sequences can ensure an enriched repertoire of highly divergent gene families.

**Fig 2.**
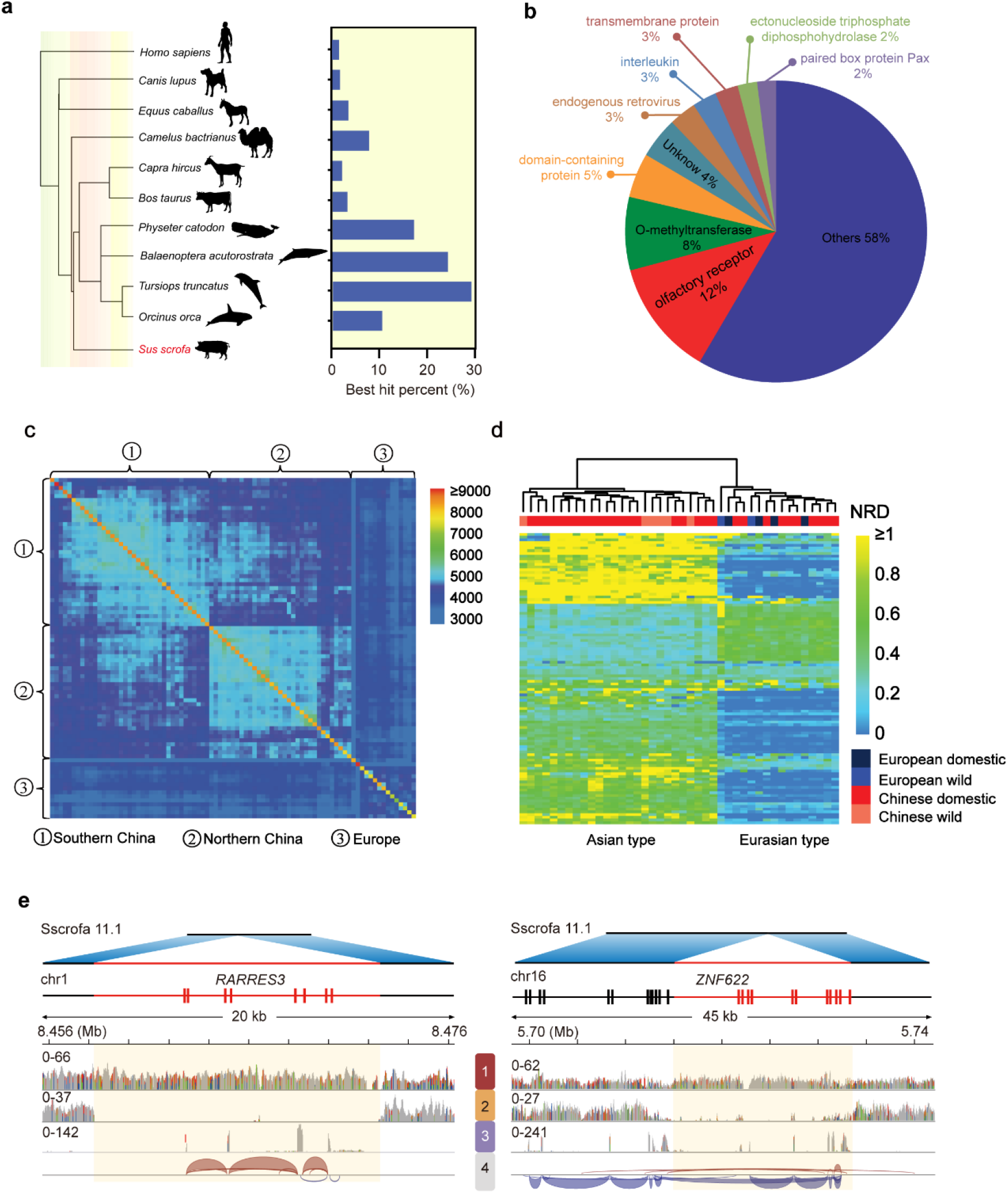
Pan-sequences validation and population-specific pattern. **a** Homolog identification of pan-sequences in ten mammalian genomes. Only the best hit was retained for each pan-sequence. **b** Number of hits in pan-sequences to Refseq protein families. **c** An 87 × 87 matrix showing the number of shared pan-sequences among all the individuals by pairs. Each cell represents the number of shared pan-sequences by two individuals. See Supplemental table 8 for the classification of each group. **d** The normalized read depth (NRD) of male-specific pan-sequences in each male. See Supplemental table 8 for the classification of each group. (Only the sequences with the frequency ranging from 0.5 to 0.9 were shown.) **e** Genes contained in the pan-sequences. One pan-sequence of 14.3 kb harbour the complete genic region of RARRES3, and another covers partial genic regions of ZNF622, representing a new splicing event. The four tracks at the bottom represent the reads mapping of whole-genome resequencing data of two samples (1–2) and the inferred exons as well as their splicing isoforms based on RNA-seq (3–4).

To explore whether these pan-sequences exhibited population-specific characteristics, we retrieved 87 publicly available pig genomes (>10× coverage) from China and Europe and aligned them to the whole pan-genome (**Supplementary Table 5**). The presence of each pan-sequence was determined by calculating their normalized read depth (NRD). The samples were clustered into three distinct groups corresponding to their geographical origin: southern Chinese, northern Chinese and European pigs (**Fig. 2c and Supplementary Table 5**), which was consist with the previously reported genetic architecture of domestic pigs (Ai, et al. 2015) and showed the presence or absence of pan-sequences can reflect the local adaptation and domestic history of pigs.

Furthermore, among the 87 pig genomes, 42 represented males, enabling us to determine the male specific sequences (**Supplementary Table 5**). We identified 10.43 Mb of male-specific sequences that were present in more than 90% of male individuals but absent in females (**Supplementary Table 6**). The distribution pattern of these male-specific pan-sequences revealed an Asian type which is confined to Asia and a Eurasian type which is distributed across Eurasia. This divergent distribution is concordant with the history of the male-biased migration from non-Asian to Asian (Guirao-Rico, et al. 2018) (**Fig. 2d**).

To determine the potential genomic positions of these pan-sequences, we aligned these pan-sequences to Sscrofa11.1 using their flanking regions. Only 19.00% (7,554) of the pan-sequences could be anchored in this way (**Supplementary Table 7**). By providing pan-sequences with refined positions, the genomic annotation could be enriched and improved. For instance, a pan-sequence of 14.3 kb containing the complete genic region of *RARRES3* is absent in Sscrofa11.1, which can be acted as a tumour suppressor or growth regulator (Shyu, et al. 2003). We further validated this gene by resequencing and RNA-seq data and found that this gene has high abundance in multiple tissues (FPKM > 1) (**Fig. 2e**). We also found that the expression of this gene is enriched in Chinese pigs, which might be a population-specific gene involved in Chinese pig growth. Another pan-sequence of 18.6 kb included six coding exons of *ZNF622*, which is also missed in Sscrofa11.1 (**Fig. 2e**). *ZNF622* played a role in embryonic development by activating the DNA-bound *MYBL2* transcription factor (Arumemi, et al. 2013). These absent six exons resulted in another spliced transcript isoform, which can be validated using the RNA-seq data.

### Constructing a more comprehensive sequence map for genomic and transcriptomic analysis

Compared with Sscrofa11.1, the mapping rate of resequencing data in the pan-genome was increased by 0.29-0.43% (**Fig. 3a and Supplementary Table 8**). Meanwhile, the mapping rate of Sscrofa11.1 in the pan-genome was decreased by approximately 1.43%, indicating that many reads had been adjusted to better positions in the pan-sequences accompanied with improved quality (**Fig. 3a, b**). The adjustment of many reads from Sscrofa11.1 to pan-sequences will result in better SNP calling efficacy. An average of 41,729 heterozygous SNPs per sample were depressed and the read depth was also adjusted to the whole-genome average level in these regions where spurious SNPs were removed (**Fig. 3b, c, d, Supplementary Fig. 4 and Supplementary Table 9**). Furthermore, 12,888 novel SNPs per individual were recovered using the pan-genome and thus provided enhanced resolution for genetic diversity studies. In addition, the mapping quality and mapping rate of RNA-seq data were also improved based on 92 samples (**Supplementary Figs. 5 and 6**). In total, 897 sequences containing 1163 potential transcripts showed appreciable expression (FPKM ≥ 1 in at least one sample). To further assess the protein-coding potential of pan-sequences, a total of 607 out of the 897 pan-sequences were predicted to have coding potential by Coding Potential Calculator (CPC) (Kong, et al. 2007). For the gene expression of pan-sequences, more expression differences were found among tissues (Pearson’s r = 0.84) than within tissues (Pearson’s r = 0.91), consistent with previous findings (Tang, et al. 2017) (**Fig. 3e and Supplementary Fig. 7**).

**Fig. 3.**
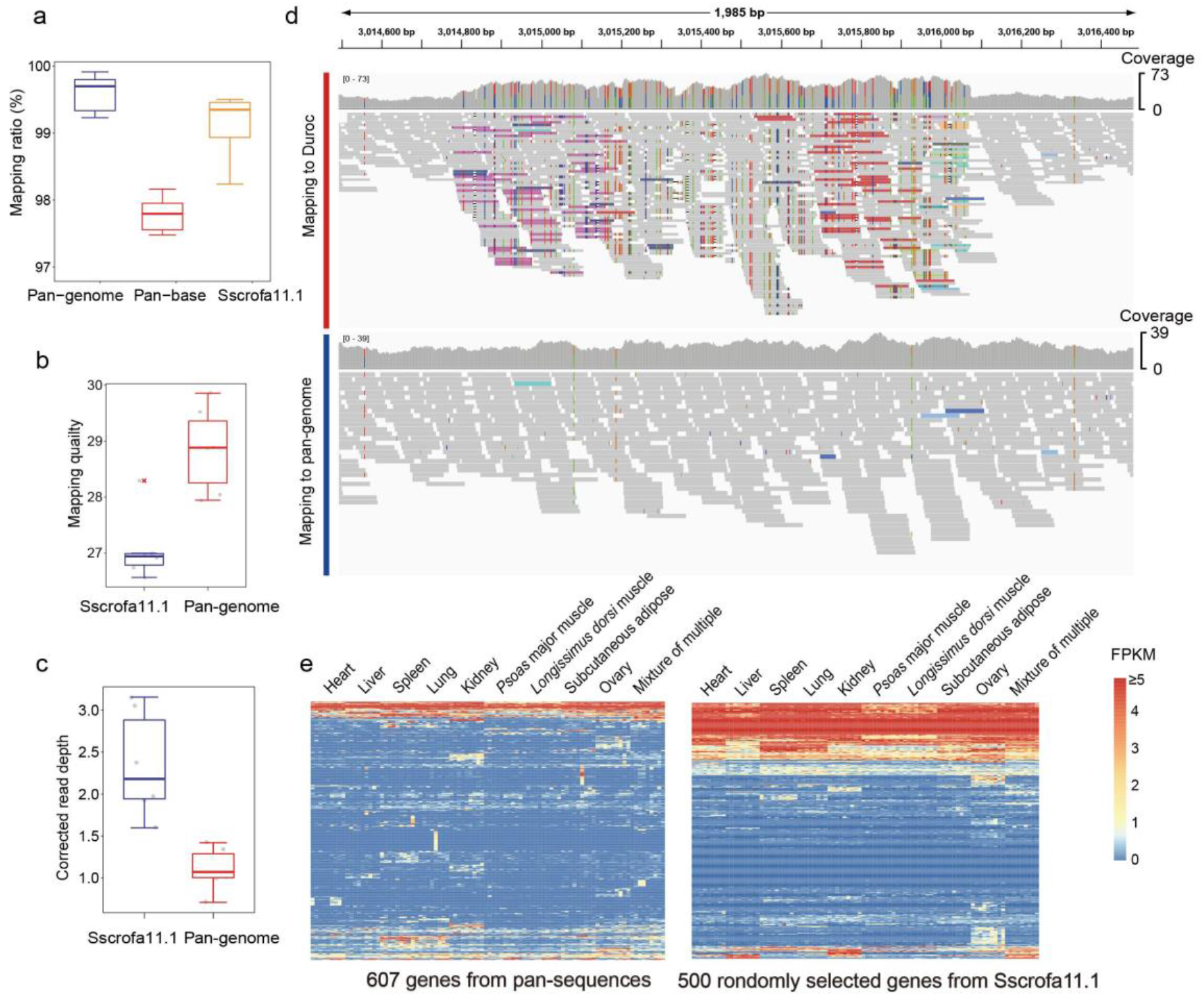
Improvements of genomic and transcriptomic analyses by using the pan-genome. **a** Comparison of the mapping ratio of resequencing data using the pan-genome versus Sscrofa11.1. **b** Comparison of read-mapping quality using the pan-genome versus Sscrofa11.1. **c** Comparison of corrected read-mapping depth using the pan-genome versus Sscrofa11.1. **d** Improved read mapping using the pan-genome versus Sscrofa11.1 as viewed with IGV. **e** Transcriptional potential of the pan-sequences.

### Hi-C based analysis revealing the characteristics of pan-sequences regarding 3D structures and their potential function

Adjacent loci generally demonstrated frequent interaction which can be determined by high-throughput Hi-C analysis (Dong, et al. 2017), thus enabling us to anchor these pan-sequences to Sscrofa11.1. Here, we generated 12 Hi-C data from 10 individuals to anchor these pan-sequences to Sscrofa11.1 by inferring their special interactions with adjacent regions (**Supplementary Tables 10 and 11**).

To evaluate the robustness and accuracy of Hi-C based localization, we comprehensively investigated the anchored results from five samples digested by the MboI enzyme and another seven samples digested with HindIII enzyme (see methods). The result indicates that the Hi-C-based localization determined by different samples has high consistency (**Supplementary Fig. 8 and 9**). A total of 7,554 sequences (19.0%) can be anchored to Sscrofa11.1 by flanking sequences. Using Hi-C based approach, another 3,447 sequences can be further anchored. Based on this result, at 100-kb resolution, we found the pan-sequences are uniformly distributed in A/B compartment which are generally euchromatic and heterochromatic regions, respectively (Dogan and Liu 2018) (**Fig. 4b**). At 20-kb resolution, we found that pan-sequences were more enriched at TAD boundary regions (**Fig. 4c**). Notably, we found that genomic variations (SNPs and CNVs) occurred more frequently at TAD boundary regions than at the TAD interior regions (**Supplementary Figs. 10, 11 and 12**), indicating that the occurrence of pan-sequences could be associated with genomic variations.

**Fig. 4.**
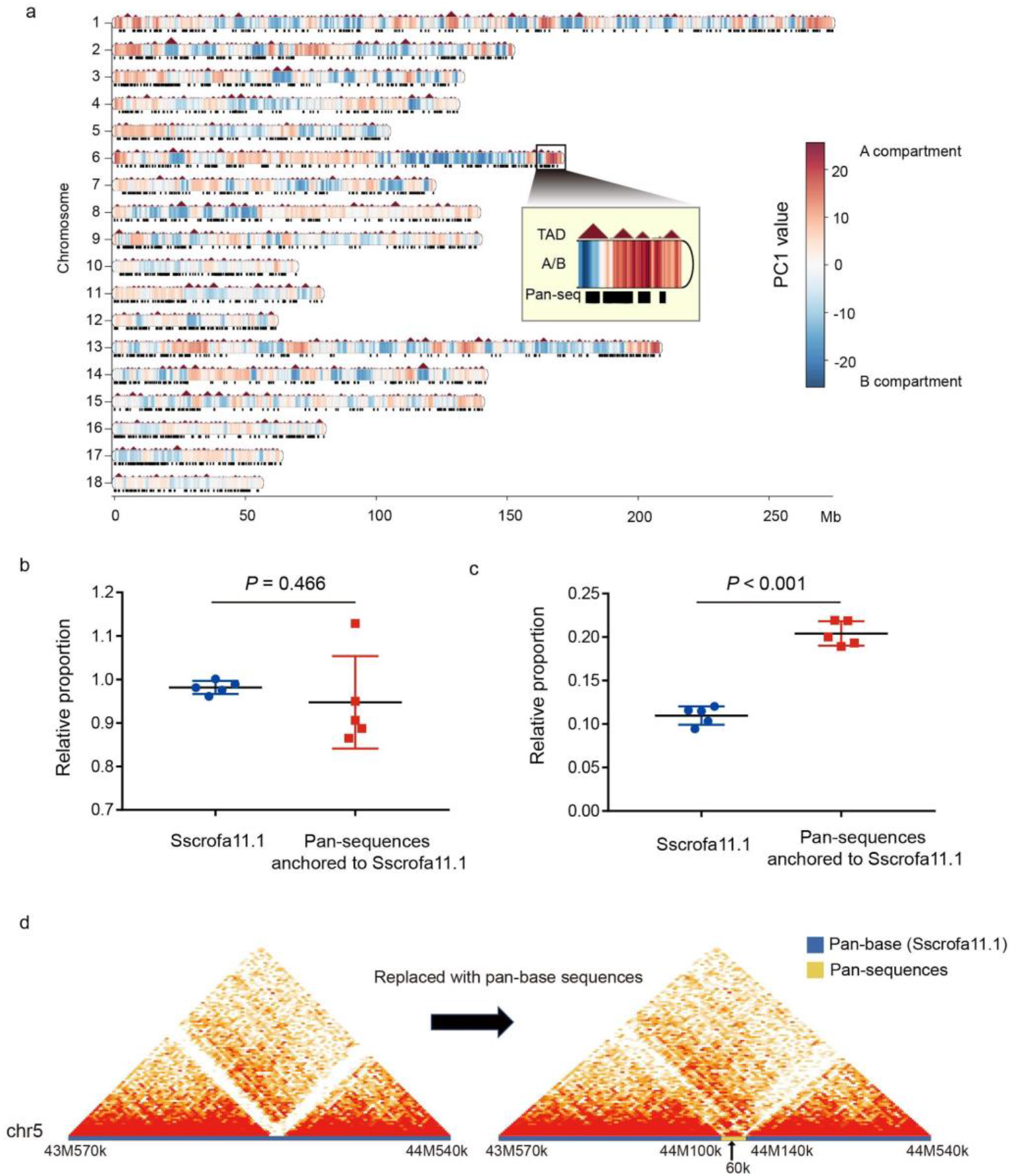
The 3D spatial structure of the pan-genome. **a** The distributions of the A/B compartment, TAD and anchored pan-sequences. **b** The relative proportion of A compartment over B compartment in length in the pig genome (left), and the relative proportion of pan-sequences located in A compartment over those located in B compartment in length (right). **c** The relative proportion of TAD boundary regions over TAD interior regions in length in Sscrofa11.1 (left) and the relative proportion of pan-sequences located in TAD boundary regions over TAD interior regions in length (right). **d** An example of improving a 3D spatial structure after replacing the weakly interacting sequences with the non-reference pan-sequences.

Based on the high genome coverage sample (∼300×), we identified 201 (5.4%) pan-sequences which were anchored to genomic regions harbouring putative enhancer elements. Furthermore, 47 pan-sequences were shown to contain enhancers by demonstrating enhancer-promoter interactions with high confidence (**Supplementary Fig. 13**). These genes which are influenced by remote regulation were significantly enriched in retinol metabolism, olfactory transduction, arachidonic acid metabolism and fatty acid degradation (**Supplementary Table 12**). When the corresponding regions of low interaction were replaced with pan-sequences, we found that 3D spatial structure was greatly improved (**Fig. 4d**). Thus, replacement with pan-sequences will help to depict the 3D structures of the whole genome.

### Pig pan-genome database

To facilitate the use of the pig pan-genome by the scientific community, a pig pan-genome database (PIGPAN) was constructed. PIGPAN is a comprehensive repository of integrated genomics, transcriptomics and regulatory data. The system diagram is shown in **Fig. 5a**. In our local UCSC Genome Browser server (Gbrowse), 17 tracks were released against the pig pan-genome (**Fig. 5b**). Users can search the database using a gene symbol or chromosome location to obtain results in terms of four aspects: (i) the reference genome and pan-sequence annotations, (ii) the gene expression in 20 corresponding tissues, seven types of regulation signals (**Supplementary Table 13**) and the conserved elements of a 20-way mammalian alignment, (iii) the chromosome localization of pan-sequences, and (iv) the haploid copy number of 87 pigs. We also provided basic search functions to retrieve basic gene information, GO annotation and KEGG pathways. Here, we present one case using PIGPAN showing the copy number difference of KIT between European and Chinese pigs (**Fig. 5c**). Moreover, users can download data from http://animal.nwsuaf.edu.cn/panPig/download.php. As the functions and associated traits of more genes in the pig genome are determined in the future, our browser will be updated regularly to meet the various needs of the scientific community.

**Fig. 5.**
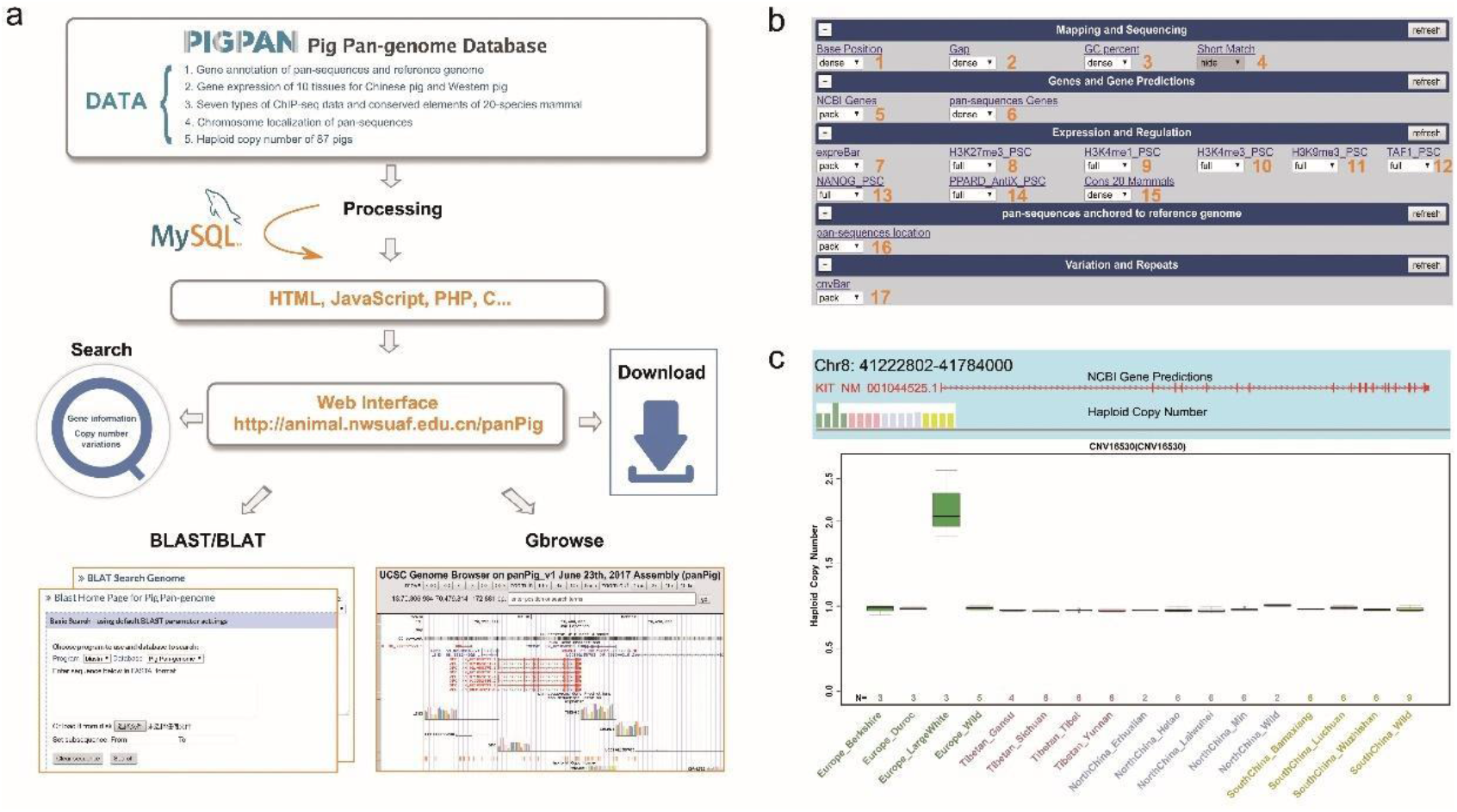
The processing pipeline used to construct PIGPAN. PIGPAN integrated genomics, transcriptomics and regulatory data. Users can search a gene symbol or a genomic region to obtain results in the form of an interactive table and graph.

## Discussion

In this study, we utilized 12 independent *de novo* assemblies (Fang, et al. 2012; Li, et al. 2013; Li, et al. 2017) and a large amount of whole-genome resequencing data to build a sequence map of the pig pan-genome. The *de novo* assemblies in the present study cover a wide range of diverse breeds across Eurasia and thus ensure a comprehensive discovery of the missing pan-sequences. These pan-sequences as well as the accompanied genomic variation and expression information will be a valuable resource for depicting the complete genetic makeup of porcine phenotypic and genomic diversity.

With the rapid decrease in the cost of generating high-quality *de novo* assemblies, the pan-genome strategy is becoming increasingly affordable and will soon become applicable for many other animal species. The importance of pan-genomes has been widely accepted in the field of plant genomics (Hirsch, et al. 2014; Schatz, et al. 2014; Golicz, Bayer, et al. 2016; Sun, et al. 2017; Zhao, et al. 2018). The high genomic plasticity of plants can result in the complete gain/loss of a large number of genes within a species (Golicz, Bayer, et al. 2016; Gordon, et al. 2017; Zhao, et al. 2018). In contrast, animal genomes are much more conserved and have longer genes with complex splicing events, which means that, generally, only intergenic or fragmented genic regions are involved in the gain/loss of genomic sequences in animals. Nonetheless, this difference does not mean that animal pan-genomes are less important. For instance, the pan-sequences that we recovered demonstrated population-specific patterns, indicating that they are potentially associated with adaptations to various environmental conditions (Li, et al. 2010; Gordon, et al. 2017). Furthermore, our research suggests that these pan-sequences may act as enhancers of some genes that regulate metabolic activity in different breeds. We also found a large number of SNPs residing in the pan-sequences, which can lead to an accurate assessment of true variations, thereby providing enhanced resolution for genetic diversity of different pig populations. The enriched genomic sequence repertoire can help in identifying causal mutations that were previously unrecognized by linkage, association and copy-number-variation studies.

In conclusion, our study has shown that the pan-genome, when used as a reference, can ensure a more comprehensive repertoire of genomic variations and can facilitate downstream genomic, transcriptomic and even 3D genome analyses. Therefore, we highlight the transition from the current reference genome to the pan-genome.

## Acknowledgements

This work was supported by research grants from the National Natural Science Foundation of China (No. 31572381) to Y.J and the Science & Technology Support Program of Sichuan (2016NYZ0042 and 2017NZDZX0002) to M.Z.L. We thank the High Performance Computing platform of Northwest A&F University for their assistance with the computing.

## Author contributions

Y.J. and M.Z. L. conceived the project and designed the research. X.T., Y.L and M.L. analysed the Hi-C data. X.T., R.L., W.F., M.L. and D.D. performed the analysis. X.T., R.L. and W.F. wrote the manuscript. Y.J, M.Z.L., X.W. revised the manuscript.

## Methods

### Construction of the pan-genome

We downloaded the publicly available pig genome assemblies of ten female and one male individuals from 11 diverse breeds (five originated in Europe and six originated in China) (**Supplementary Fig. 2 and Table 2**) (Fang, et al. 2012; Li, et al. 2013; Li, et al. 2017). To identify the sequences which cannot align to the reference genome, we split the 11 assemblies by gap region and iteratively aligned them to the reference pig genome assembly (Sscrofa11.1) using the BLASTN (Camacho, et al. 2009). Sscrofa11.1 was masked by WindowMasker (Morgulis, et al. 2006) before alignment to speed up the alignment process. The sequences with <90% identity and ≥300 bp in length were retained. After that, these low-identity sequences were aligned to each other to remove redundancy. Finally, a non-redundant set of 72.5 Mb of sequences from 11 assemblies was obtained; these sequences were defined as pan-sequences.

The Duroc genome (Sscrofa11.1) plus these 72.5 Mb pan-sequences made up the pan-genome.

### Determining the characteristics of the pan-sequences

To explore whether the pan-sequences have homologous regions across species and are potential to be functional, we aligned these sequences to ten mammalian reference genomes (i.e., *Homo sapiens*, *Camelus bactrianus*, *Equus caballus*, *Canis lupus familiaris*, *Capra hircus*, *Bos Taurus*, *Orcinus orca*, *Physeter catodon*, *Balaenoptera acutorostrata scammoni*, *Tursiops truncates*) to search for any matches (*E*-value ≤ 1e-5) using BLASTN (Camacho, et al. 2009) (**Supplementary Table 4**). Only the best hit was remained for each query.

To validate the authenticity of these pan-sequences and identify assembly-specific sequences, we aligned all of them to the each of the 11 *de novo* pig assemblies to search for any matches (≥90% coverage and ≥95% identity) using BLASTN (Camacho, et al. 2009). If the sequence of an assembly does not have a high similarity with other assemblies, this sequence is considered as the assembly-specific sequences.

### Population-based resequencing and CNV calling

We downloaded the whole genome resequencing data for 71 domestic pigs and 16 wild boars for population analysis of pan-sequences. The sequences data were retrieved from NCBI under the Bioproject PRJNA213179, PRJNA281548, PRJNA309108 and PRJEB9922 (Ai, et al. 2015; Frantz, et al. 2015; Jeong, et al. 2015; Li, et al. 2017) (**Supplementary Table5**). After alignment using BWA (version 0.7.15-r1140) (Li and Durbin 2009) with default parameters, we used CNVcaller (Wang, et al. 2017) to calculate the normalized read depth (NRD) of each sequences. The presence and absence of each pan-sequence were then determined by NRD.

### ChIP-seq short-read alignment and peak calling

To confirm the content of regulatory elements in pan-sequences, we downloaded seven ChIP-seq data from NCBI Bioproject PRJNA152995, including H3K27me3, H3K4me1, H3K4me3, H3K9me3, NANOG, PPARD AntiX and TAF1 signals (**table S13**) (Xiao, et al. 2012). Sequencing reads were aligned to pig pan-genome using BWA (version 0.7.17-r1188) (Li and Durbin 2009) with default parameters. Low-quality and multiple-mapping reads were removed using SAMtools (Li, et al. 2009) with option “-q 20”. Enriched regions (or peaks) were called (p < 1e-5; no filtering on fold enrichment or FDR correction) using MACS (version 2.1.1) (Zhang, et al. 2008) with total DNA input as control.

### Identification of male-specific sequences

There are 42 males and 45 females in our whole genome resequencing data (**Supplementary Table 8**). We compared the normalized read depth (NRD) between females (NRD < 0.1, sample size = 42) and males (0.2 < NRD < 0.7, sample size = 45) to identify the putative male-specific pan-sequences. Thus, we identified 1,638 male-specific scaffolds (**Table S9**) which were present in most all of male individuals (frequency ≥ 50%) but absent in females (frequency = 0) with a combined length of 10,432,972bp (**Supplementary Table 6**).

### Gene annotation and functional enrichment analysis

Homology-based and *de novo* prediction were used to annotate protein-coding genes. For homology-based prediction, pan-sequences were aligned onto the repeat-masked assembly using TblastN (Camacho, et al. 2009) with an *E*-value cutoff of 1e-5. Aligned sequences as well as corresponding query proteins were then filtered and passed to GeneWise to search for accurately spliced alignments (Doerks, et al. 2002). For *de novo* prediction, GenScan (Burge and Karlin 1998), Augustus (Stanke, et al. 2006), and geneid (Blanco, et al. 2007) were then used to predict genes.

Annotated genes of novel sequences were analysed for Kyoto Encyclopedia of Genes and Genomes (KEGG) terms and pathway enrichment using KOBAS (Xie, et al. 2011).

### SNP calling

To verify whether using the pan-genome as reference could improve SNP calling efficacy, we randomly selected six pig samples (ranging from 10 to 30× coverage) (**Supplementary Table 8**) and mapped their clean reads to the pan-genome and Sscrofa11.1 for comparison. Duplicate reads were removed using Picard Tools. Then, the Genome Analysis Toolkit (GATK, version 3.6) (McKenna, et al. 2010) was used to detect SNPs. The following criteria were applied to all SNPs: (1) Variant confidence/quality by depth (QD) > 2; (2) RMS mapping quality (MQ) > 30.0.

### RNA-seq analysis and noncoding RNA prediction

The 92 strand-specific RNA-seq data (7-10 tissue libraries for each of 10 individuals) were downloaded from the NCBI database (Bioproject: PRJNA311523) (Li, et al. 2017). All reads were mapped to the pan-genome by HISAT2 (Kim, et al. 2015). Transcripts including novel splice variants were assembled using StringTie version 1.2.2 (Pertea, et al. 2015) and the FPKM (Fragments Per Kilobase per Million mapped reads) values for these transcripts and genes in each sample were determined using Ballgown (Frazee, et al. 2015). Finally, transcripts with FPKM ≥ 1 in at least one sample were retained. After assembling and quantifying all transcripts, the transcripts of pan-sequences were used for identification of high confidence coding RNA by Coding Potential Calculator (CPC) (Kong, et al. 2007) online.

### Materials for Hi-C experiment

Liver of BH-33, BH-34, BH-35, and BH-36 were collected from four female 2-years-old Bama minipigs. Liver of F2 were collected from a 90-days-old female fetus of Bama minipig. Ear skin fibroblasts DB-2 and DB-3 were established by using two 12-days-old female Large White pigs. Ear skin fibroblasts XYZ were established by using a 2-years-old female Wild Boar. Embryonic fibroblasts RC-7 and RC-8 were established by using two 40-days-old female fetus of Chinese Rong Chang pig. Mature adipocytes DB-2-Y and DB-3-Y were derived from pre-adipocytes which were established by using the same pigs of Ear skin fibroblasts DB-2 and DB-3, by inducing adipogenic differentiation.

All of the fibroblasts were grown in DMEM Dulbecco’s Modified Eagle Medium (DMEM, 11995-065, Gibco) containing 10% Fetal Bovine Serum (FBS, 10099-141, Gibco) and 1× penicillin/streptomycin (P/S,15140-122, Gibco), incubated at 37°C in 5% CO2.

Pre-adipocytes were cultured in 10%FBS/DMEM-F12 (11330-032, Gibco) with 1×P/S until confluence and induced to differentiation as previously described. Briefly, two days’ post-confluence, cells were exposed to differentiation medium containing 0.5 mmol/L isobutylmethylxanthine (I5879, Sigma), 1 μmol/L dexamethasone (D2915, Sigma), 850 nmol/L insulin (I6634, Sigma), 1 μmol/L rosiglitazone (R2408, Sigma) and 10% FBS for three days. At the end of day 3, the differentiation medium was replaced into maintenance medium with only 850 nmol/L insulin, 1 μmol/L rosiglitazone and 10% FBS, and replenished every other day. After the differentiation process, at least 90% of the cells had accumulated lipid droplets at day 15, and were used as mature adipocytes (DB-2-Y and DB-3-Y).

### Hi-C experimental method

Hi-C experiment on cells were performed according to the previously published Hi-C protocol with some minor modifications (Lieberman-Aiden, et al. 2009).
Briefly, 25 million (M) cells were resuspended in 45 ml serum free DMEM, and 37% formaldehyde was added to obtain a final concentration of 2% for chromatin cross-linking. Cells were incubated at room temperature (20–25 °C) for 5 minutes, then glycine was added to obtain a final concentration of 0.25 mol/L to quench the formaldehyde. The mixture was incubated at room temperature for 5 minutes, and subsequently on ice for at least 15 minutes. Fixed cells were lysed using a Dounce homogenizer in the presence of cold lysis buffer (10 mmol/L Tris-HCl, pH 8.0, 10 mmol/L NaCl, 0.2% IGEPAL CA-630, and 1× protease inhibitor solution). Chromatin digestion (restriction enzyme HindIII), labelling, and ligation steps were performed according to the original protocol (Lieberman-Aiden, et al. 2009). After deproteinization, removal of biotinylated free-ends, and DNA purification, Hi-C libraries were controlled for quality and sequenced on an Illumina Hiseq X Ten sequencer (paired-end sequencing with 150 bp in read length).

Hi-C experiment on liver tissue were performed as previously described using the MboI restriction enzyme (Rao, et al. 2014), with minor modifications pertaining to handling flash frozen primary tissues (Leung, et al. 2015). Briefly, 0.5 g flash frozen liver tissue was pulverized in liquid nitrogen. Then cross-linking by 37% formaldehyde in a final concentration of 4% and incubated at room temperature for 30 mins. Glycine was added to obtain a final concentration of 0.25 mol/L to quench the formaldehyde. The mixture was incubated at room temperature for 5 minutes, and subsequently on ice for at least 15 minutes. Cross-linked liver cells were filtered through 70-μm and 40-μm nylon cell strainers and spinning down to collect the liver cells. About 25 mg liver cell precipitate was used for Hi-C library preparation. The Hi-C library preparation procedure was performed as previously described using the MboI restriction enzyme (Rao, et al. 2014).

### Hi-C reads mapping, filtering, and generation of contact matrices

Pre-processing paired-end sequencing data, reads mapping as well as filtering of mapped di-tags was performed using the Juicer pipeline (v.1.8.9) (Durand, et al. 2016). Briefly, short reads were mapped to pan-genome using BWA (version 0.7.15-r1140) (Li and Durbin 2009). Reads of low mapping quality were filtered using Juicer with default parameters, discarding the invalid self-ligated and un-ligated fragments, as well as PCR artefacts. Filtered di-tags were further processed with Juicer command line tools to bin di-tags (10 kb bins) and to normalize matrices with KR normalization (Knight and Ruiz 2013). Valid Hi-C read pairs should harbour more intrachromosomal (cis) interactions than inter-(trans) (**Supplementary Table 11**). To improve resolution, we combined the Hi-C data from the same tissue of same pig breed after we randomly extracted 20 Gb data for correlation coefficient test. We combined Hi-C data from DB-2 and DB-3 (Pearson’s r = 0.99); RC-7 and RC-8 (Pearson’s r = 0.99); DB-2-Y and DB-3-Y (Pearson’s r = 0.96). After combined samples, all processes were done in all the data. Normalized interaction matrices were generated at two resolutions of low (100 kb) and high (20 kb) respectively (**Supplementary Figure 14**).

### Identification of compartment A and B

Identification of compartment A/B was performed as previously described using the 100-kb interaction matrix (Lieberman-Aiden, et al. 2009). Principal component analysis (PCA) was performed to generate the first principal component (PC1) vectors of each chromosome, and Spearman’s correlation between PC1 and genomic characteristics (gene density and GC content) were then calculated. GC content (%) for each bin (100-kb bin sizes) was calculated using SeqKit (v.0.8.0) (Shen, et al. 2016). Gene density (number of genes per bin) was calculated based on the number of promoters [from −2,000 to +500 bp of transcription start site (TSS)] located in (namely more than 50% of the region should be overlapped) each bin. Compartment A and B were determined by the PC1 values. Bins with positive Spearman’s correlation between PC1 values and genomic features were assigned as compartment A, otherwise B.

### Identification of topologically associating domains (TADs) and topological boundaries

Higher-resolution TAD calls were generated following the previously described procedure by using the directionality index (DI) metric (Dixon, et al. 2012). DI was calculated using raw interaction counts between 20-kb bins to capture observed upstream or downstream interaction bias of genomic regions. A hidden Markov model (HMM) was then used to predict the states of DI for final TAD generation. The same criteria 400 kb (distance between two adjacent TADs) was used to distinguish unorganized chromatin from topological boundaries. That is, the topological boundaries are less than 400 kb and unorganized chromatin is larger than 400 kb.

### Locating pan-sequences on Sscrofa11.1 based on Hi-C

We normalized all Hi-C matrices on the same scale by KR normalization (Knight and Ruiz 2013), ensuring that any differences between Hi-C are not attributable to variation in sequence length. The maximum 100-kb bin of each pan-sequence interacted (Interaction intensity ≥ 5) was collected as the potential location of pan-sequences. Starting with the filtered 100-kb resolution bin of pan-sequences, we get the higher resolution interval of 20 kb by taking the maximus 20-kb bin with each 100-kb bin.

### Identification of putative promoter and enhancer interactions

We kept the interactions identified by PHYCHIC (Ron, et al. 2017) with FDR < 0.01 as high confidence interactions and used them to identify promoter-enhancer interactions (PEI). Promoter segment was determined as a region from −2,000 to +500 bp of the transcription start site (TSS). When at least half of a promoter segment was in either one of the two bins which involved in a chromatin interaction, this interaction was defined as a putative promoter interaction.

The bins which are distal (at least 40 kb upstream or downstream) from the promoter and demonstrate the strongest interaction with the promoter than other regions were determined as the enhancer interacting with the corresponding promoter. This interaction of the two bins corresponding to the promoter and enhancer was defined as a potential PEI. If our pan-sequences were located on a bin harbouring an enhancer of a PEI, the pan-sequences could be potentially involved in the regulatory functions of the enhancer. If the pan-sequences further demonstrate interactions with the promoter of the same PEI, the involvement of the pan-sequences in the regulatory functions of the enhancer would be regarded as highly confident and the pan-sequences could be a potential enhancer itself.

### The pig pan-genome web server

The web interface of PIGPAN was built by combining Apache web server, PHP, HTML, JavaScript and relational database MySQL. Users can use all online resources without preregistration. Our browser can be divided into two parts: frontend and backend interfaces. The frontend consists of a home page, a download page and several search pages. The MySQL relational database server stores 16 tables including gap information, GC percent, seven regulatory signals of potential stem cells (H3K27me3, H3K4me1, H3K4me3, H3K9me3, NANOG, PPARD AntiX, TAF1), conserved elements of 20-species mammal, haploid copy number of 87 pigs, gene expression, location of pan-sequences and gene annotation. The appropriate index was built on the corresponding retrieval columns of the table. When a user submits an entry, the backend will respond quickly to execute an SQL statement. PHP and JavaScript manage the data analysis processes and visualize the results. Moreover, we have introduced web-based software such as BLAST (Camacho, et al. 2009), BLAT (Kent 2002) and Gbrowse (Casper, et al. 2018). Accordingly, users can query data with rapid visualization in Gbrowse or enter a query sequence to search for homologous regions in the genome. PIGPAN was tested in all major modern internet browsers, including Firefox, Chrome, Internet Explorer, Safari and Opera. Therefore, PIGPAN is a robust and easy-to-use website to facilitate the search for and visualization of results for pig pan-genome analyses.

## Data availability

The sequencing reads of each sequencing library have been deposited at NCBI for Hi-C data (Project ID: PRJNA496307). The assembly of pig pan-genome and subsequent analysis results are available in our PIGPAN website
(http://animal.nwsuaf.edu.cn/code/index.php/panPig). All other data supporting the findings of this study are available in the article and its supplementary information files or are available from the corresponding author on request.

